# Effects of the Salinity under Soilless Culture Systems on Gamma Linolenic Acid Levels in Borage Seed Oil

**DOI:** 10.1101/454405

**Authors:** Miguel Urrestarazu, Victor Manuel Gallegos-Cedillo, Francisca Ferrón-Carrillo, José Luis Guil-Guerrero, Teresa Lao, Juan Eugenio Álvaro

## Abstract

Borage is a well-known plant of great importance in human nutrition and health. Expanding knowledge of particular plants that have anti-cancer products is a global concern. There is substantial information regarding the benefits, presence and extraction of gamma linolenic acid (GLA) in different plants around the world, especially in borage seeds. However, there is little information concerning the effects of the salinity of the nutrient solution on the growth and presence of GLA in borage seeds. The objective of this work was to determine the optimal salinity of the nutrient solution for obtaining GLA in soilless cultivation systems. Borage plants were grown in coconut fibre and provided three treatments of nutrient solution of 2.20, 3.35 and 4.50 dS m^−1^, increasing solution salinity with the standard nutrient solution of concentrated macronutrients as a reference. Vegetative growth, seed production and GLA ratio were measured. The results of vegetative development and GLA production doubled and tripled with the increase in salinity of the nutrient solution, respectively.

## Introduction

Borage (*Borago officinalis* L.) is a native plant of the Mediterranean region that is currently cultivated around the world to produce its seed oil. The quantity of borage seed marketed each year is variable, fluctuating between 500 and 2000 t worldwide. The global borage oil market exceeded 1,500 t in 2015. It is expected that the borage oil market will have an estimated value of 54.9 million dollars by 2024 [1, 2]. According to the Ministry of Agriculture and Fisheries, Food and Environment (MAPAMA) Spain reported 810 ha with an output of 14,001 t in 2016, with fresh consumption being the main destination of production followed by animal and human consumption. Further, the main producing provinces were Navarra and Aragón (81.88%), La Rioja (16.53%) and Madrid (1.57%) [3].

At present, it is well-established that there is a growing interest in producing not only foods with high organoleptic qualities but also functional foods [4]. Many recent therapeutic and preventive medicines include the use of traditional plant-based preparations [5]. Borage seed oil has been used as a treatment for various degenerative diseases [6, 7]; more recently, the supplementation of gamma linolenic acid (GLA) from borage seed oil has been shown to protect DNA by modulating oxidative genetic damage in *Drosophila melanogaster* [8].

The effects of salinity on general productivity have been well-established [9]. There is rich information regarding the effects of the salinity of nutrient solutions on the nutrient composition of many crops, such as tomatoes [4, 10, 11]. However, there is very little information regarding the effects of salinity on the composition of GLA in borage seed oil.

The objective of this work was to determine the effects of the salinity in the nutrient solution on the productivity of borage crops and the presence of fatty acids in their seeds.

## Materials and methods

### Plant growth conditions

The study was performed at the University of Almeria, Spain, in a Raspa y amagado greenhouse similar to that described by [12]. The vegetal material used was borage (*B. officinalis* L.), transplanted in a state of 4 true leaves in 20 L containers filled with coconut fibre substrate that was composed of 85% fibre and 15% peat, whose physical characteristics were described by [13]. A planting density of 1.25 plants m^−2^ was used. The transplant period was from August 15, 2016 to July 31, 2017. The average temperature in the greenhouse at night was 15-20 °C and 20-35 °C during the day without supplemental lighting.

### Treatments

The plants were fertigated daily with different salinity levels in the nutrient solution. The treatments were 2.20, 3.35 and 4.50 (dS m^−1^) of electrical conductivity (EC) of the nutrient solution, based on [14] (Table 1). Concentrated mother solutions were used for the macronutrients until the desired EC was reached, and the corresponding proportion of micronutrients was subsequently added. The pH of the nutrient solutions was always maintained at 5.8 with the addition of nitric acid.

**Table 1.** Nutrient solution compositions for different salinities.

| EC<br>(dS m<sup>-1</sup>) | pH | Macronutrient (mM) |  |  |  |  |  | Micronutrient (μM) |  |  |  |  |  |
| --- | --- | --- | --- | --- | --- | --- | --- | --- | --- | --- | --- | --- | --- |
|  |  | NO<sub>3</sub><sup>-</sup> | H<sub>2</sub>PO<sub>4</sub><sup>-</sup> | SO<sub>4</sub><sup>2-</sup> | K<sup>+</sup> | Ca<sup>2+</sup> | Mg<sup>2+</sup> | Fe | Mn | Cu | Zn | B | Mo |
| 2.20<sup>1</sup> | 5.80 | 10.25 | 1.50 | 1.75 | 4.75 | 5.00 | 1.51 | 15 | 10 | 0.75 | 5 | 30 | 0.5 |
| 3.35 | 5.80 | 12.81 | 1.88 | 2.19 | 5.95 | 6.25 | 1.89 | 15 | 10 | 0.75 | 5 | 30 | 0.5 |
| 4.50 | 5.80 | 15.38 | 2.25 | 2.63 | 7.13 | 7.50 | 2.27 | 15 | 10 | 0.75 | 5 | 30 | 0.5 |

<sup>1</sup>Based on [14].

The EC of 2.20 dS m^−1^ was considered the standard saline treatment.

### Fertigation system

Three drainage collection trays and control drippers were installed per treatment. To adjust the treatments, in each irrigation the volume (L), pH and EC (dS m^−1^) of the supplied irrigation and drainage were measured; the volume was measured with a graduated cylinder with precision to the hundredths, and pH and EC were measured with an MM40^+^(Hach^®^ LPV2500.98.0002, Spain). Each new irrigation was performed when 10% of the water readily available in the substrate had been used, and the volume needed to obtain between 20-30% of drainage was added [13, 15, 16] while removing the pegs to avoid the preferential distribution channels of the nutrient solution [17].

### Growth parameters

The evaluation of growth parameters was 181 days after transplantation. The experimental unit was four plants per repetition and four repetitions per treatment. The parameters measured were plant height (cm), number of leaves plant^−1^, leaf thickness (measured with a micrometric screw in the midpart of the margin while avoiding the ribs), stem diameter (cm; measured with a digital calliper (Stainless Hardened, Spain)), root length (cm; measured with a tape measure), and leaf area (m^2^ plant^−1^; measured with an AM350 Area Meter (ADC BioScientific Ltd., Hertfordshire, United Kingdom)). The plants were divided by their different organs; the fresh weight of roots, stems and leaves was obtained, then the dry weight was obtained by placing the material in an oven (Thermo Scientific Heratherm^®^, Germany) at 75 ºC until achieving a constant weight. A precision analytical balance (Adventurer^®^ Analytical OHAUS Modelo AX 124/E, USA) was used, expressing the result as g plant^−1^.

### Estimation of daily flower number

To estimate the number of flowers per day, all the flowers that opened each day were identified with a label indicating the date and treatment. This procedure was repeated for six fortnights. The number of flowers that opened daily per plant and treatment were recorded.

### Pollination and seed production

The pollination of the flowers was accomplished manually with the help of a brush from 8 to 10 am, when the flowers entered anthesis.

Harvesting was performed manually when the seeds reached physiological maturity (fruit dehiscence and dark-coloured seeds), to have the highest seed quality [18, 19, 20]. Immediately after harvest, the seeds were placed in a glass desiccator (Vacuumfest DURAN^®^, Germany) with 1-3 mm of silica gel for storage until measurement in the laboratory. Subsequently, the width and length (mm) were measured for 400 seeds per treatment using a digital calliper (Stainless Hardened, Spain). Similarly, the total number of seeds per plant^−1^ and fortnight^−1^ were obtained; in addition, the dry seed weight for each treatment was determined with a precision analytical balance (Adventurer^®^ Analytical OHAUS Model AX 124/E, USA).

The harvest index was calculated from the total dry weight (g plant^−1^) and the seed production (g plant^−1^), expressed as a percentage.

### Oil extraction and transesterification

Extraction and trans-esterification were performed simultaneously, and fatty acid (FA) analyses and quality control were carried out according to previous reports [21, 22].

Seeds were ground in the lab with the aid of a mortar and pestle, and then 150–200 mg was taken for direct methylation and further Gas-Liquid Chromatography (GLC) analyses. Each sample was analysed in triplicate. Ground seeds were weighed in 10 mL test tubes, and then 1 mL of the methylation mixture (methanol:acetyl chloride 20:1 v/v) and 1 mL of *n*-hexane were added. Tubes were capped and later heated at 100 °C for 30 min. Afterwards, the tubes were cooled to room temperature, 1 mL of distilled water was added, and after centrifugation (3,500 rpm, 3 min), the upper hexane layer was removed for GLC analysis [23].

### Fatty acid analyses

Fatty acid methyl esters (FAME) were analysed using a Focus GLC (Thermo Electron, Cambridge, UK) equipped with a flame ionization detector (FID) and an Omegawax 250 capillary column (30 m x 0.25 mm i.d. x 0.25 µm film thickness; Supelco, Bellefonte, USA). The temperature program was as follows: 1 min at 90 °C, heating until 220 °C at a rate of 10 °C min^−1^, maintenance at a temperature of 220 °C (2 min), then heating until 250 °C at a rate of 10 °C min^−1^ and then maintenance at a constant temperature of 250 °C (1 min). The injector temperature was 250 °C with a split ratio of 50:1. The injection volume was 4 µL. The detector temperature was 260 °C. Nitrogen was used as the carrier gas (1 mL min^−1^). Peaks were identified by retention times obtained for known FAME standards (PUFA N^o^. 1, 47033; methyl γ-linolenate 98.5% purity, L6503; and methyl stearidonate 97% purity, 43959 FLUKA) from Sigma, (St. Louis, USA), and FA levels were estimated using methyl pentadecanoate (15:0; 99.5% purity; 76560 Fluka) from Sigma as an internal standard [24].

### Design and analysis of experiments

The experimental design was a randomized complete block system with 3 treatments and 4 repetitions (n=4). The experimental unit consisted of 4 plants [25]. The results of the agronomic variables were subjected to analysis of variance and Tukey’s test (p ≤ 0.05 was considered significant). The processing of the data was done using Statgraphics Centurion^®^ XVI. II.

## Results and discussion

### Vegetative growth

Table 2 shows that the means of the different recorded vegetative growth parameters were similar to other borage crops grown in open air [1, 26]. In the control treatment, the nutrient solution at the EC standard of 2.20 dS m^−1^ showed an average total fresh weight greater than conventional borage crops reported by several previous studies, such as [27].

**Table 2.**
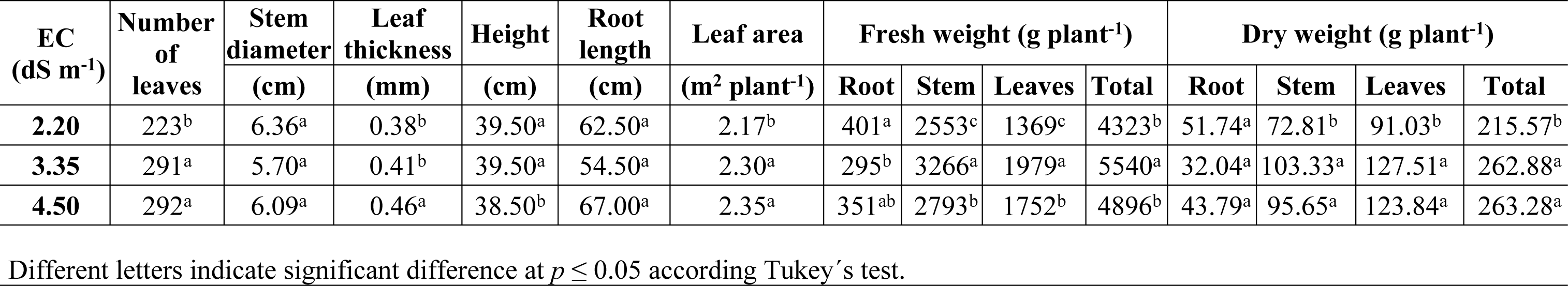
Growth parameters of borage crop (*B. officinalis* L.) versus electric conductivity (EC) of nutrient solution.

The electrical conductivity of the nutrient solution showed an important effect on vegetative development (Table 2). While the development of the root was not clearly and significantly affected, shoots increased from an average of 6% in the leaf area to 30% in the fresh or dry weight of the leaves when the nutrient solution EC increased from 2.20 to 3.35 dS m^−1^. When the EC increased from 3.35 to 4.50, the fresh (but not dry) weight of the leaves decreased by 12%.

Jaffel, Sai [28], who increased the EC from a standard nutrient solution with NaCl, recorded similar results with low salinity (25 mM), while they reported that a 5 dS m^−1^ increase above this EC resulted in a significant decrease in the vegetative growth of leaves, stems, roots and buds. In contrast, these same authors [29] recorded a significant reduction in the production of biomass from a nutrient solution with the addition of 25 mM NaCl, as also recorded by [30] from an EC of 5 dS m^−1^.

#### Flower and seed yield

The highest ECs showed a much higher precocity in the first fortnight (Fig 1). In contrast, [28] significantly reduced flowering from the very first salinity treatment applied (25 mM NaCl). Over the complete crop cycle, the number of flowers at the highest EC was significantly higher. The number of viable seeds showed similar behaviour. Our EC control treatment (2.20 dS m^−1^) showed a much higher seed productivity (13.97 g plant^−1^) than the non-saline treatment recorded by [29] (1.15 g plant^−1^) (Table 3). The highest EC treatments, 3.35 and 4.50 dS m^−1^, generated significant increases of 27 and 40% higher than the previous EC level, respectively.

**Fig 1.**
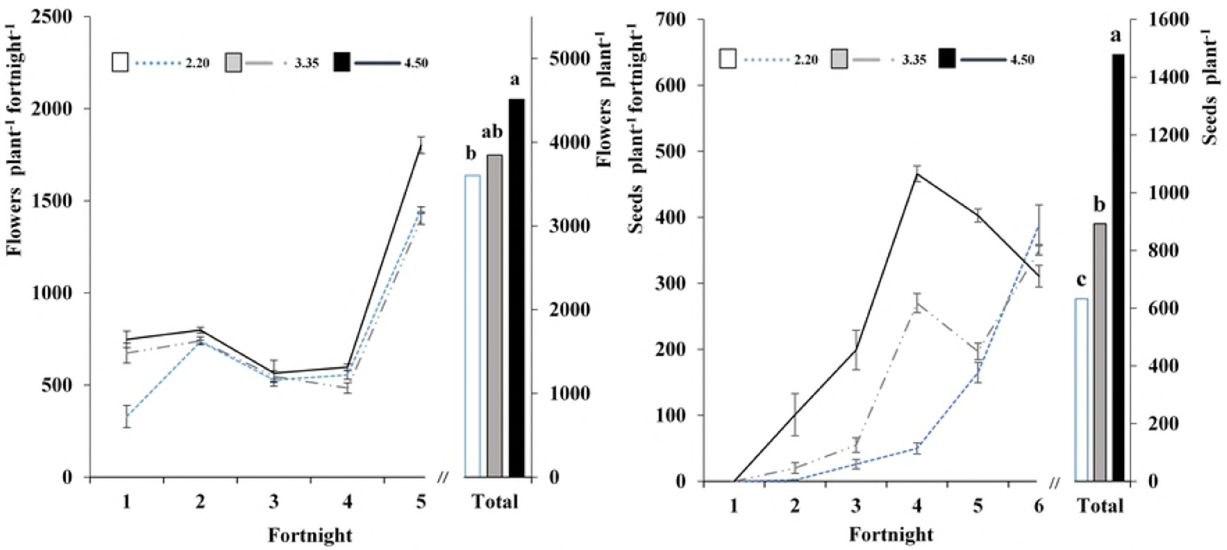
Flowers and seeds of borage crop versus electric conductivity (dS m^−1^). Different letters indicate significant difference at p ≤ 0.05 according to Tukey’s test.

**Table 3.** Seed parameters and harvest index of borage crop (*B. officinalis* L.) versus electric conductivity (EC) of nutrient solution.

| EC (dS m<sup>-1</sup>) | Seed<br>(g plant<sup>-1</sup>) | Weight of<br>1000 seeds (g) | Length | Width | Harvest index |
| --- | --- | --- | --- | --- | --- |
|  |  |  | (mm) |  |  |
| 2.20 | 13.97<sup>c</sup> | 22.1<sup>a</sup> | 5.29<sup>a</sup> | 3.09<sup>a</sup> | 0.06<sup>b</sup> |
| 3.35 | 19.27<sup>b</sup> | 21.6<sup>a</sup> | 5.13<sup>a</sup> | 3.08<sup>a</sup> | 0.07<sup>b</sup> |
| 4.50 | 31.67<sup>a</sup> | 21.4<sup>a</sup> | 5.39<sup>a</sup> | 3.12<sup>a</sup> | 0.12<sup>a</sup> |

Different letters indicate significant difference at $p \leq 0.05$ according to Tukey's test.

Similarly, a significant doubling of the number of seeds was recorded in the EC treatment of 4.50 dS m^−1^ relative to the lowest EC. The 3.35 dS m^−1^ treatment resulted in an intermediate seed number, but that number was also significantly higher than the lowest EC treatment. These positive results are clearly contrary to those obtained by [29], who, by increasing the EC of the nutrient solution by means of NaCl (of similar EC salinity to our treatment of 3.35 dS m^−1^), reported substantially decreased seed production and did not obtain many seeds with their treatments of higher salinities (similar to or greater than our treatment of 4.50 dS m^−1^). This difference in results could be justified because it is well known that a greater benefit is achieved in productivity - at equal ECs of the nutrient solution - when these are obtained with a proportional increase of the macronutrients compared to when NaCl is added [31, 32, 33].

The unit weight of the seeds was significantly higher than those obtained by previous authors, such as [1], and similar to the slightly below average weight reported by [34]; the height and width were slightly higher than the dimensions described in Flora Ibérica (2012) by [35]. EC treatments did not affect either the weight of the seeds or their size.

Harvest index median values were similar to those reported by authors such as [36], but the highest EC increased notably and significantly compared to the lower EC treatments.

#### Fatty acid production

Table 4 shows the FA composition of and production by borage. The average FA ratios were similar to borage plants collected from the Maghreb [21], Spain and Sardinia [24], Tunisia [29], and Chile [1].

**Table 4.** Fatty acid (FA) levels in borage (*B. officinalis* L.) crop seeds for different nutrient solution electrical conductivity (EC) values.

| EC<br>(dS m<sup>-1</sup>) | Fatty acids (FA% of total FAs) |  |  |  |  |  |  |  |  |  |  |  |  |  | FA amount | FA | 18:3n6 |
| --- | --- | --- | --- | --- | --- | --- | --- | --- | --- | --- | --- | --- | --- | --- | --- | --- | --- |
|  | 12:0 | 14:0 | 16:0 | 16:1n7 | 18:0 | 18:1n9 | 18:2n6 | 18:3n6 | 18:3n3 | 20:0 | 20:1n9 | 22:0 | 22:1n9 | 24:1n9 | (g 100 g<sup>-1</sup><br>seed) | g m<sup>-2</sup> | g m<sup>-2</sup> |
| **2.20** | 0.11 | 0.09<sup>b</sup> | 12.94<sup>a</sup> | 0.27<sup>a</sup> | 4.11<sup>b</sup> | 22.3<sup>a</sup> | 34.2<sup>a</sup> | 17.9<sup>c</sup> | 0.23<sup>a</sup> | 0.27<sup>b</sup> | 3.48<sup>b</sup> | 0.17<sup>c</sup> | 2.25<sup>b</sup> | 1.33<sup>b</sup> | 31.7<sup>b</sup> | 5.52<sup>c</sup> | 0.99<sup>c</sup> |
| **3.35** | - | 0.09<sup>b</sup> | 12.81<sup>a</sup> | 0.26<sup>a</sup> | 4.40<sup>a</sup> | 21.6<sup>a</sup> | 33.2<sup>a</sup> | 18.7<sup>b</sup> | 0.22<sup>a</sup> | 0.32<sup>a</sup> | 3.73<sup>a</sup> | 0.20<sup>b</sup> | 2.71<sup>a</sup> | 1.51<sup>a</sup> | 32.6<sup>a</sup> | 7.83<sup>b</sup> | 1.47<sup>b</sup> |
| **4.50** | - | 0.11<sup>a</sup> | 11.96<sup>b</sup> | 0.23<sup>b</sup> | 4.15<sup>b</sup> | 18.9<sup>b</sup> | 35.4<sup>a</sup> | 20.0<sup>a</sup> | 0.22<sup>a</sup> | 0.28<sup>ba</sup> | 3.79<sup>a</sup> | 0.26<sup>a</sup> | 2.92<sup>a</sup> | 1.60<sup>a</sup> | 32.5<sup>a</sup> | 12.84<sup>a</sup> | 2.57<sup>a</sup> |

Different letters indicate significant differences at $p \leq 0.05$ according to Tukey's test. (n =4)

The GLA production of the control treatment was 0.99 g m*^−2^*, which was much higher than the 0.72 g m*^−2^* reported by [27], most likely due to the better development conditions that are obtained with a soilless cultivation system and greenhouse conditions. The increase in salinity exerted a significant and beneficial effect both on the general concentration of FAs and on those most beneficial for human health. Both the total production of FA and GLA were practically tripled at ECs from 2.20 to 4.50 dS m^−1^. Jaffel, Sai [28] also found that some degree of salinity in the borage crop increased the metabolic activity of important reactive oxygen-scavenging enzymes, such as superoxide dismutase, and had no induction of activity of catalase—an ascorbate peroxidase—and a slight increase in glutathione reductase activity.

## Conclusion

An increase in the nutrient solution from 2.20 to 4.50 dS m^−1^ through a balanced ratio of macronutrients provides an elevated and significant increase in vegetative growth. With an increase in EC up to 4.50 dS m^−1^, floral and seed production doubled compared to an EC standard of 2.20 dS m^−1^. The ratio of fatty acids and gamma-linolenic acid doubled or tripled with a salinity of 4.50 dS m^−1^.

## Acknowledgment

This work was supported by the Spanish Ministry of Education and Science Project (FEDER AGL2015-67528-R).

The authors gratefully acknowledge the support of the Mexican National Council for Science and Technology (CONACYT) for its financial support of this work.

